# NGOMICS-WF, a Bioinformatic Workflow Tool for Batch Omics Data Analysis

**DOI:** 10.1101/475699

**Authors:** Weizhong Li

**Affiliations:** J Craig Venter Institute, La Jolla, California, United States of America

## Abstract

In recent years, high throughput next generation omics data have been broadly used in many research fields. However, analyzing omics data is still very challenging, especially for researchers with less experience in bioinformatics, software, and programming. Omics data analysis workflows usually are composed of many processes involving different software tools that demand different compute resources. So, it is often difficult to perform a full workflow in an automated way, despite that many computational tools and resources are available. Here I present NGOMICS-WF, a lightweight workflow tool for batch omics data analysis. This workflow tool has been used in several projects. It can assist researchers to configure and run workflows for various types of omics data on Linux computers or Linux computer clusters. The software package is available from https://github.com/weizhongli/ngomicswf. In addition to the workflow tool, several pre-configured workflows that were tested and published are also available in the package. These pre-configured workflows can be directly used or be used as templates to configure new workflows to analyze metagenomic, metatranscriptomic, RNA-seq and 16S data.

## 1 Introduction

In recent years, the extraordinary developments of experimental techniques, especially the newer generation of sequencing technologies, have enabled more omics-based studies at a bigger scale. The type of omics data has also expended remarkably. Data analysis of omics-based project has been becoming more demanding ever since. Along with the development of experimental techniques, computational tools, databases, resources and analysis methods have also been rapidly evolving. For many common omics-based projects, data processing and analysis usually require many different software tools, making it extremely difficult to perform end-to-end analysis in an automated way and to scale up to large number of samples.

A key to address these issues is through a workflow or a pipeline. Workflow and pipeline have some distinct meanings in some aspects, however, in the area of computational processing of omics data, workflow and pipeline don’t have clear boundaries and they have been used interchangeably by the communities. There are some existing workflow software tools in the past and many novel tools were developed recently. Many workflow tools have common features for workflow management and pipeline integration, such as handling dependency between processes, automation of full workflow run, scaling up with multiple input datasets, workflow post-error restart, real-time monitoring of process and so on. Traditional workflows, such as Taverna [1] and Kepler [2], can handle very complicated processes and offer a graphic user interface for workflow implementation. Workflow functionality by Galaxy [3] provides web-based workflow development and execution. Some other workflow tools, without a graphic interface, are also very popular due to easy usage. Examples of them include Snakemake [4], a system that uses rules similar to UNIX command “make” for dependency management. As a general-purpose workflow tool, Luigi (https://github.com/spotify/luigi) is also widely used in bioinformatic pipelines. Features and aspects for many existing workflow tools were summarized in a recent review [5].

The workflow tool presented here is a lightweight tool, which was originally developed from 2010 and has been used in our group for many years in projects involving analysis of genomic, metagenomic, transcriptomic data [6]. This tool is also behind several web servers developed by our group [7, 8]. This tool was implemented to facilitate bioinformatics developers to configure and run a pipeline on Linux cluster or standalone Linux system. The software and several pre-configured workflows that have been tested and published are available from https://github.com/weizhongli/ngomicswf as open source software.

## 2 Results

### 2.1 Workflow system

NGOMICS-WF is different from other workflow tools in design strategy. We use a straightforward template-based method for users to configure a workflow. This allows unrestricted flexibility in workflow management. The structure of NGOMICS-WF is illustrated in Fig. 1. In summary, the workflow system includes the following key components:

**Figure 1.**
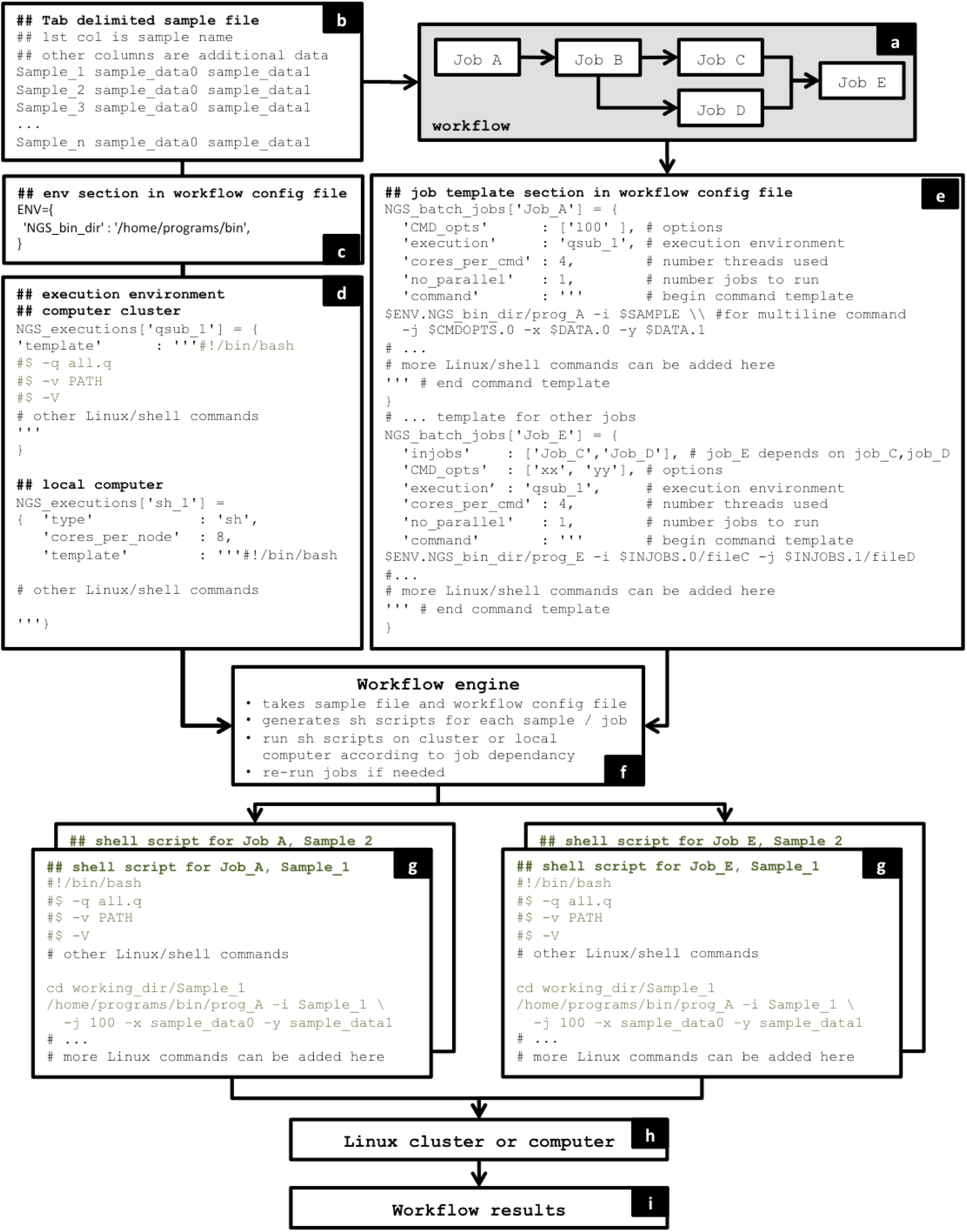
Schema of NGOMICS-WF

- **Workflow:** this is the conceptual workflow (Fig. 1a).
- **Workflow template**: this is the workflow configuration file made by a user. It defines the individual jobs in workflow, job dependencies, computing requirements and other parameters (Fig. 1c, 1d, 1e).
- **Workflow engine**: it is software that takes a user-configured workflow template and runs the workflow on a list of omic datasets (Fig. 1f).
- **Execution environment**: this refers to either a local computer or a computer cluster where the workflow is executed (Fig. 1d).
- **Third-party programs and data:** these are bioinformatic programs (e.g. tools for reads mapping or assembly), resources (e.g. reference databases) and data to be used in the workflow.
- **Utility scripts:** many scripts that are useful for handling sequence files, parsing results, job partition are included in the package.

The most important part for users is the workflow template, through which the workflow is configured. This template is an editable text file composed in Python. Here, only very limited Python knowledge is needed to edit a workflow template. Each job is defined as a block in the workflow template file. This block declares various job parameter: upstream dependent jobs, compute resources to run the job (e.g. a queue of a computer cluster or the local computer), number of cores needed (in case of multithreading programs), number of computer nodes needed (in case of parallel jobs to be submitted to multiple nodes at a computer cluster), job failing test condition (e.g. if output file has zero size), and actual command lines.

The workflow template file usually also claims one or multiple execution resources, such as a local computer or one or more queues in a computer cluster. Here, key parameters such as the number of cores should be defined. The workflow template also defines local variables, such as PATH to executables or third party programs.

Given one or a list of input datasets (Fig. 1a), when a user executes a workflow, the workflow engine will convert the workflow template into shell scripts for each job for each dataset, according to a small set of pre-defined rules (Fig. 1g). These shell scripts will be executed either on a local computer or be submitted to a computer cluster when the job dependencies are met (Fig. 1h). The workflow engine will automatically run through all the jobs defined in the workflow template (Fig. 1i).

### 2.2 Major features

This system has several important features.

- **Lightweight design**: for users with working knowledge in Linux computer and cluster, this system is extremely easy to use. Although the workflow template file is written in Python, only very basic knowledge in Python is needed.
- **Flexibility**: workflow configuration is done through a text editor; users can directly configure jobs at command line level and have the flexibility to use Linux shell commands. Each job in the workflow template is basically a block of shell commands, which can be as simple as a single command, or can be multiple command lines, or even with loop or other programming structures and with internal shell variables.
- **Computing resource management**: this tool makes it straightforward to interact with job execution environments to utilize the available compute cores and memory, without over executing. The workflow engine will only submit job when there is enough computing resources.
- **Parallel job handling**: jobs that are independent will be executed in parallel as long as there are enough computing resources. For example (Fig. 1a), after job B is finished, job C and D can be performed in parallel. When multiple samples are processed, jobs from different samples will also be executed in parallel.

### 2.3 Pre-configured workflow examples

Together with the workflow engine, several pre-configured workflows are also available from the software package. These can be used directly in analyzing metagenomic, metatranscriptomic and 16S rRNA data, with minor modifications. They are also served as workflow examples for users to configure new workflows.

One workflow was based on our web server webMGA [7] and was the tool for analysis in a human microbiome project [6]. Another workflow performs OTU clustering for Miseq sequencing based 16S rRNA sequences [9].

## 3 Implementation

NGOMICS-WF was previously written in Perl and recently re-written in Python. Besides Python and its core component, which are installed by default on most Linux systems, no additional Python packages are needed. The tool works under a standalone Linux computer or cluster with a queue system. It has been tested with Open Grid Engine (OGE), previously known as Sun Grid Engine (SGE) on computer clusters. It has also been widely tested on Amazon cloud environments, where virtual computer clusters can be set up. Many other computer cluster job queuing systems have very similar set of commands as OGE, so NGOMICS-WF should work with these systems with minor modification.

The pre-configured workflows were written in Python. Many utility scripts included in NGOMICS-WF were written in Perl.

## 4 Conclusion

Here I present an effective bioinformatic workflow tool with several unique features for omics data analysis. The software package is freely available from https://github.com/weizhongli/ngomicswf. Documentation and examples are also available with the package.

## Funding

This work was partly supported by Award R01HG005978 to WL from the National Human Genome Research Institute (NHGRI). The content is solely the responsibility of the authors and does not necessarily represent the official views of the NHGRI or the National Institutes of Health. This work was partly supported by the U. S. Department of Agriculture (USDA) National Institute of Food and Agriculture under Award No. 2013-67015-22957 to WL. Names or commercial products in this publication is solely for the purpose of providing specific information and does not imply recommendation or endorsement by USDA. The funders had no role in study design, data collection, and analysis, decision to publish, or preparation of the manuscript. This work was partly supported by the J. Craig Venter Institute.

### Conflict of Interest

None declared.

